## Supplementary material for "Phenotypes associated with genes encoding drug targets are predictive of clinical trial side effects"

SUPPLEMENTARY FIGURES

Figure S1. Correlation of independent variables in side effect model. All variables are binary; correlations displayed are odds ratios (ORs); asterisks show p-value from Fisher's exact test (*, p < 0.05; **, p < 0.005; ***, p < 0.0005).

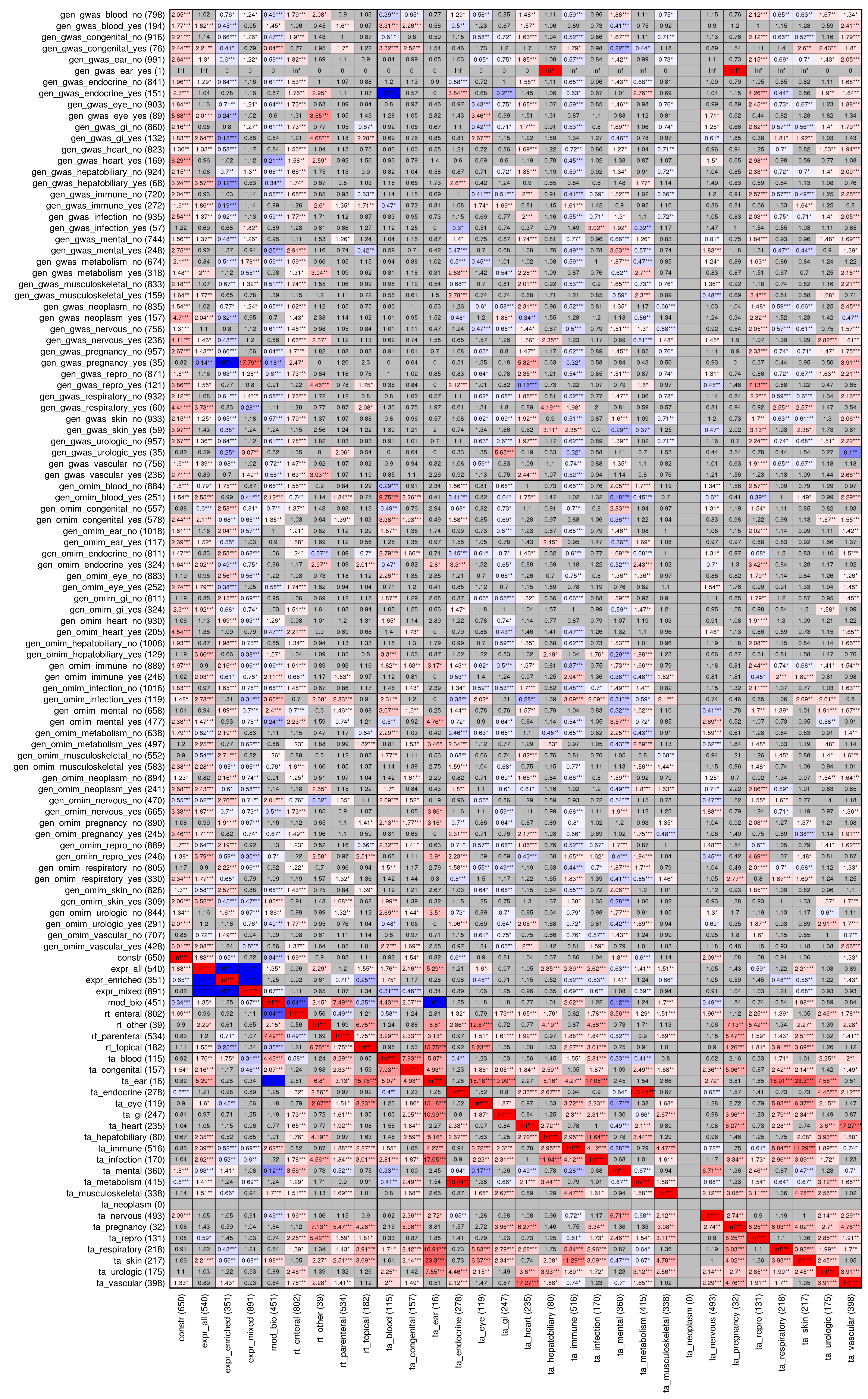

Figure S2. All coefficients from each of the 18 side effect regression models, where “on-indication” side effects are retained in the data. Asterisks show p-value of the coefficient (*, p < 0.05; **, p < 0.005; ***, p < 0.0005).

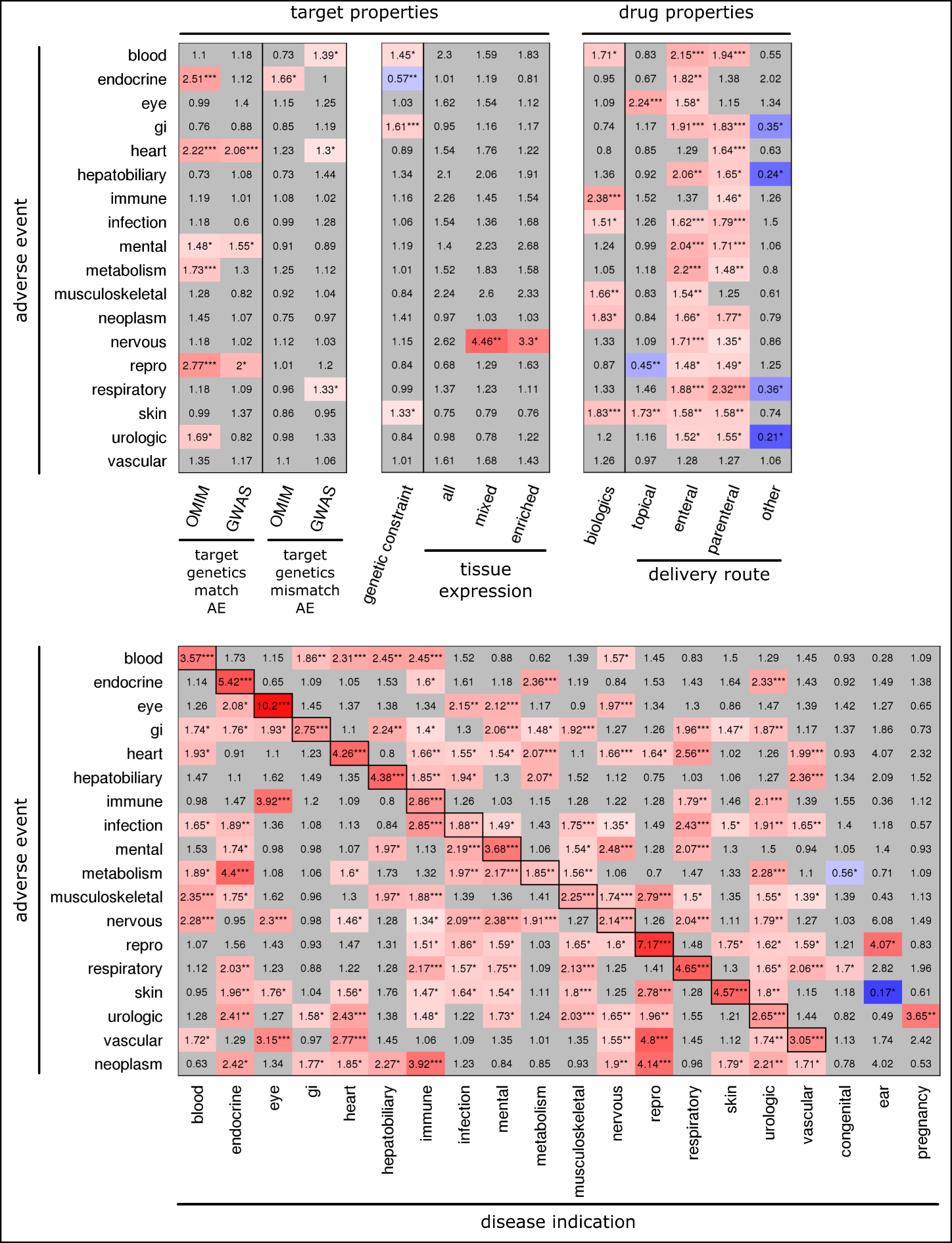

Figure S3. All coefficients from each of the 18 side effect regression models, where only “off-indication” side effects are retained in the data. Asterisks show p-value of the coefficient (*, p < 0.05; **, p < 0.005; ***, p < 0.0005).

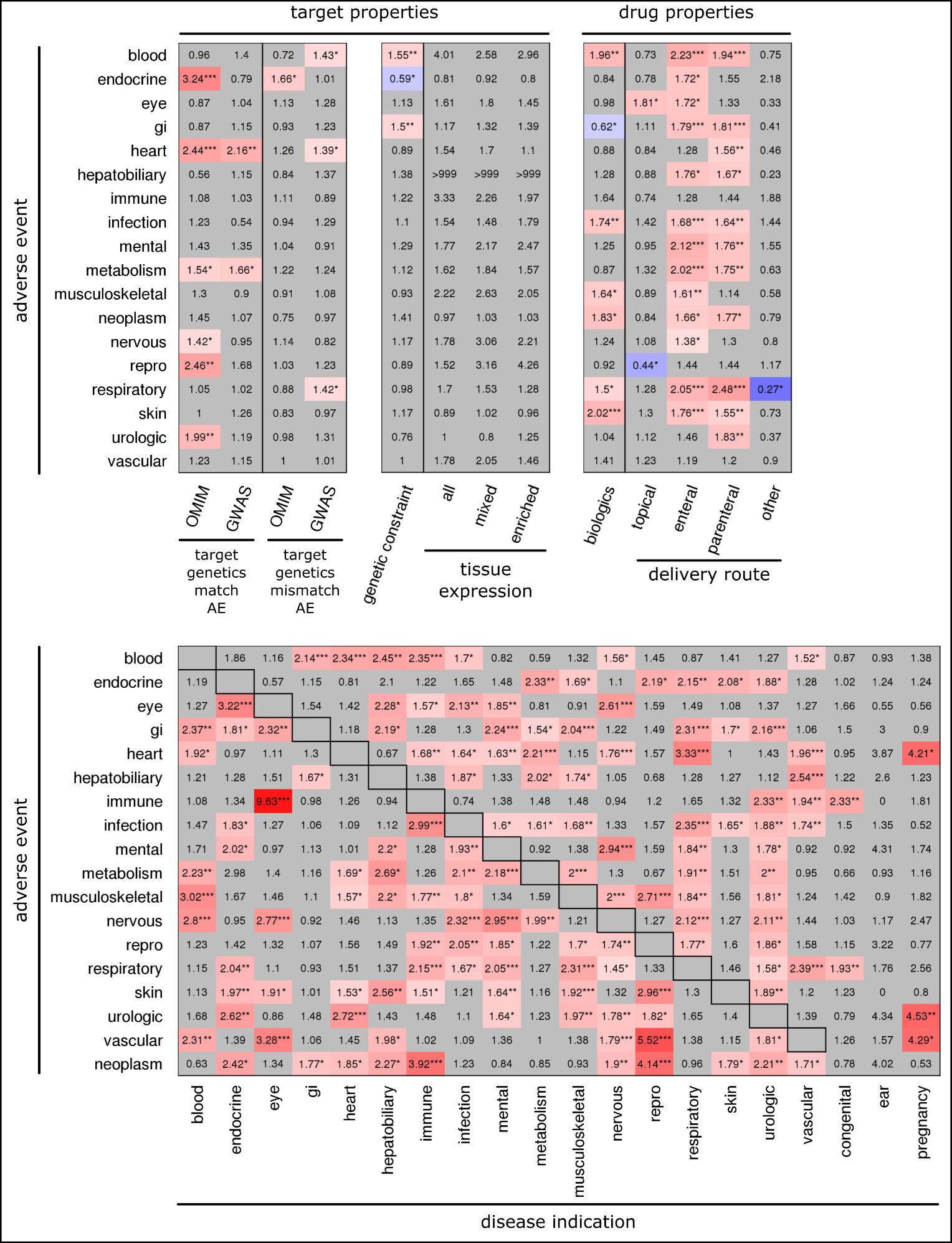

Figure S4. Comparison of the rate at which all possible drug-SE (side effect) combinations are manifested as side effects in the clinical trial data set, between drug-SE pairs with or without genetic support. OR, odds ratio; p, p-value from Fisher’s exact test (two-tailed). The top two panels show analyses based on all 38,199 drug-SE combinations; the bottom two panels show analyses based on 33,489 “off-indication” drug-SE combinations. The left two panels show analyses based on Mendelian genetic data; the right two panels show analyses based on GWAS genetic data. Error bars represent the 95% confidence interval of the reported proportions. Underlying data and confidence intervals for the OR values are shown in **Table S2**.

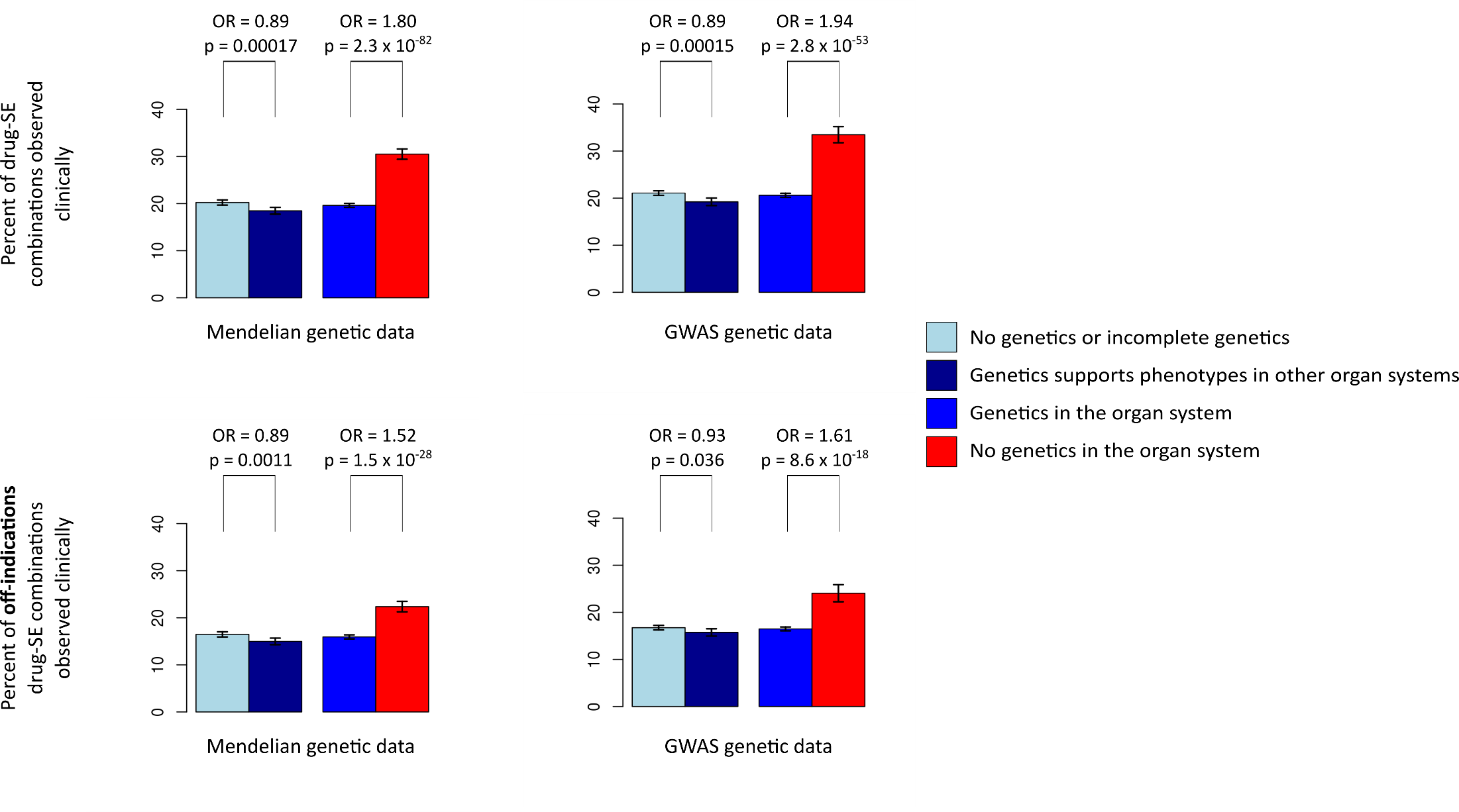

Figure S5. Summary of side effect modeling results with quantitative, tissue-specific target gene expression values included in the models. LEFT: coefficients of the Mendelian genetics predictors in each of the 18 side effect models (from Table S11). RIGHT: coefficients of the GWAS genetics predictors in each of the 18 side effect models (from Table S11). Coefficients from the regression models were exponentiated to obtain odds ratios. Along each row, circular points indicate odds ratios from the glm models; error bars are 95% confidence intervals from the glm models; large eight-pointed stars indicate odds ratios when the genetics predictors are selected as predictors in the glmnet models (which are built using feature selection, lasso regularization, and cross-validation). Small asterisks offset from the data indicate p-values of the the Fisher’s exact test (LEFT) and p-values of the coefficients from the glm models (CENTER and RIGHT); (*, p < 0.05; **, p < 0.01; ***, p < 0. 001).

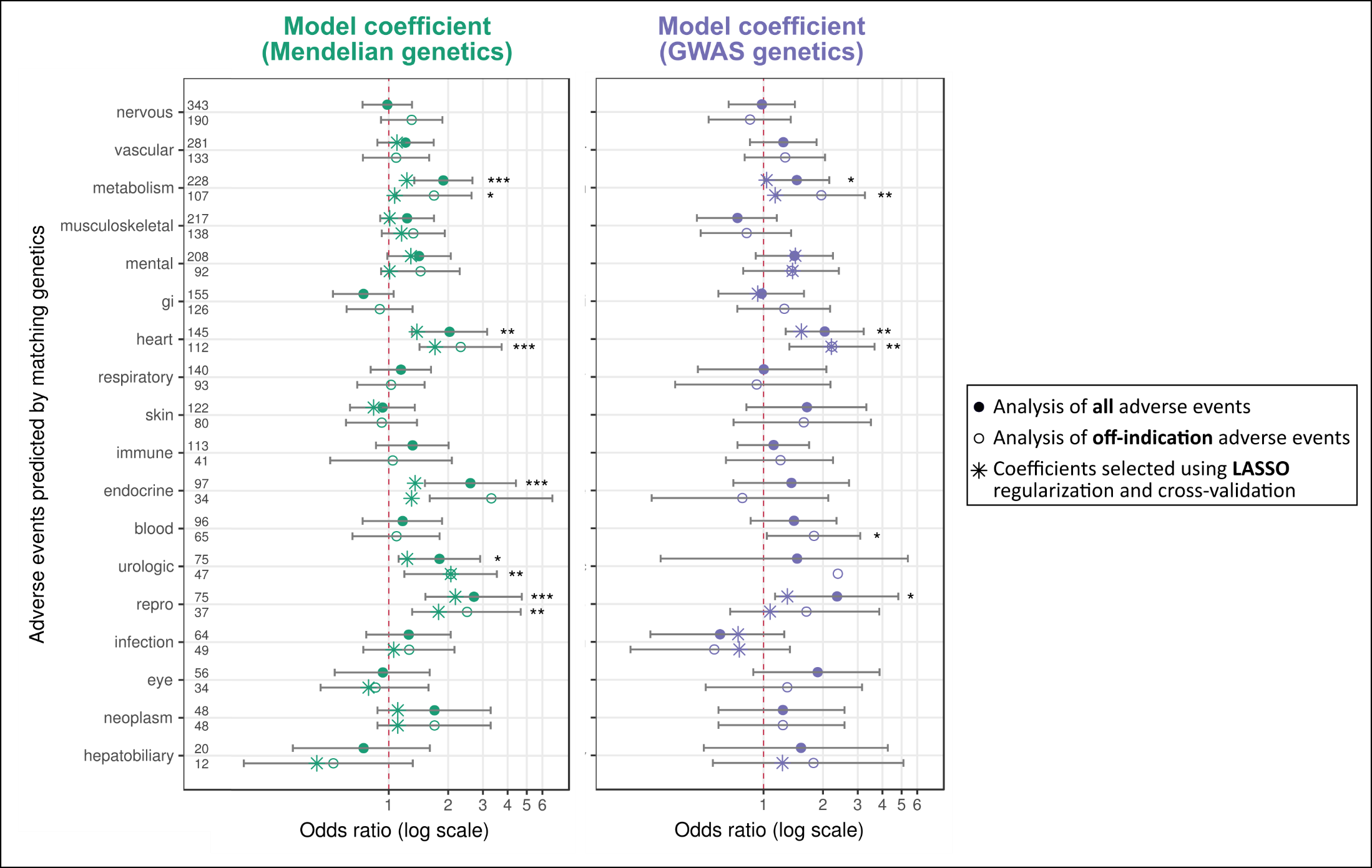
